# Transcriptome analysis reveals temporally regulated genetic networks during *Drosophila* border cell collective migration

**DOI:** 10.1101/2023.09.27.559830

**Authors:** Emily Burghardt, Jessica Rakijas, Antariksh Tyagi, Pralay Majumder, Bradley J.S.C. Olson, Jocelyn A. McDonald

## Abstract

**Background:** Collective cell migration underlies many essential processes, including sculpting organs during embryogenesis, wound healing in the adult, and metastasis of cancer cells. At mid-oogenesis, *Drosophila* border cells undergo collective migration. Border cells round up into a small group, detach from the epithelium, and migrate – at first rapidly through the surrounding tissue, then slower, with the cluster rotating several times before stopping at the oocyte. While specific genes that promote cell signaling, polarization of the cluster, formation of protrusions, and cell-cell adhesion are known to regulate border cell migration, there may be additional genes that promote these distinct dynamic phases of border cell migration. Therefore, we sought to identify genes whose expression patterns changed during border cell migration.

**Results:** We performed RNA-sequencing on border cells isolated at pre-, mid-, and late-migration stages. We report that 1,729 transcripts, in nine co-expression gene clusters, are temporally and differentially expressed across the three migration stages. Gene ontology analyses and constructed protein-protein interaction networks identified genes expected to function in collective migration, such as regulators of the cytoskeleton, adhesion, and tissue morphogenesis, but also a notable enrichment of genes involved in immune signaling, ribosome biogenesis, and stress responses. Finally, we validated the *in vivo* expression and function of a subset of identified genes in border cells.

**Conclusions:** Overall, our results identified differentially and temporally expressed genetic networks that may facilitate the efficient development and migration of border cells. The genes identified here represent a wealth of new candidates to investigate the molecular nature of dynamic collective cell migrations in developing tissues.

## Introduction

Cell migration shapes tissues throughout the life of an organism, from embryonic development to wound healing in the adult, including in various disease states such as in cancer. Cells can migrate individually, like in immune surveillance, or collectively in larger interconnected and coordinated groups of cells [1; 2; 3]. A wide variety of collective cell migrations occur *in vivo*, including *Drosophila* dorsal closure, vertebrate blood vessel formation and remodeling, vertebrate neural crest migration, wound healing, and tumor metastasis. Despite the diversity of cell types and organisms, the cellular and molecular mechanisms that govern collective cell migration are remarkably conserved. In response to external signals, cell polarization and cytoskeletal rearrangements define the collective’s leader cells, which form F-actin-rich protrusions to provide traction but also sense the extracellular environment [2; 4]. Leader and follower cells are linked through adhesion proteins and the actomyosin cytoskeleton, which together promote contraction and movement of the entire group. Because cells move inside developing tissues, organs, and organisms, our understanding of the molecular mechanisms that coordinate various collective cell migration behaviors is still incomplete.

One of the best studied genetic models of collective cell migration is the *Drosophila* border cells, which move as a small group during development of the ovary. The ovary is composed of multiple strings of progressively developing egg chambers bundled together [5]. Each egg chamber consists of an oocyte and 15 supportive germline nurse cells in the center, surrounded by a monolayer of somatic epithelial follicle cells [6]. During mid-oogenesis, four to six follicle cells at the very anterior of the egg chambers are specified to become border cells by the polar cells, a pair of non-motile follicle cells [5; 7]. Border cells then surround the polar cells to form a migratory cluster (**Figure 1A**). After assembly, the border cell cluster delaminates from the follicular epithelium and migrates between the large nurse cells (**Figure 1B**). Border cell migration is a highly dynamic and active process. Throughout their migration, border cells continuously extend and retract protrusions. There is also no fixed leader cell. Instead, individual border cells exchange places and move within the cluster [8]. Border cells stop migrating when they reach the oocyte, where they become epithelial again in a process termed “neolamination” (**Figure 1C**) [9]. After delamination, it can take border cells three to four hours to travel the entire ∼150-200 µm distance to the oocyte [8]. Border cells are then joined by the migrating centripetal cells to completely enclose the anterior side of the oocyte [10]. Eventually, border cells, polar cells, and a subset of centripetal cells form a structure in the eggshell called the micropyle, which serves as an entry point for sperm to fertilize the oocyte [11; 12; 13].

**Figure 1.**
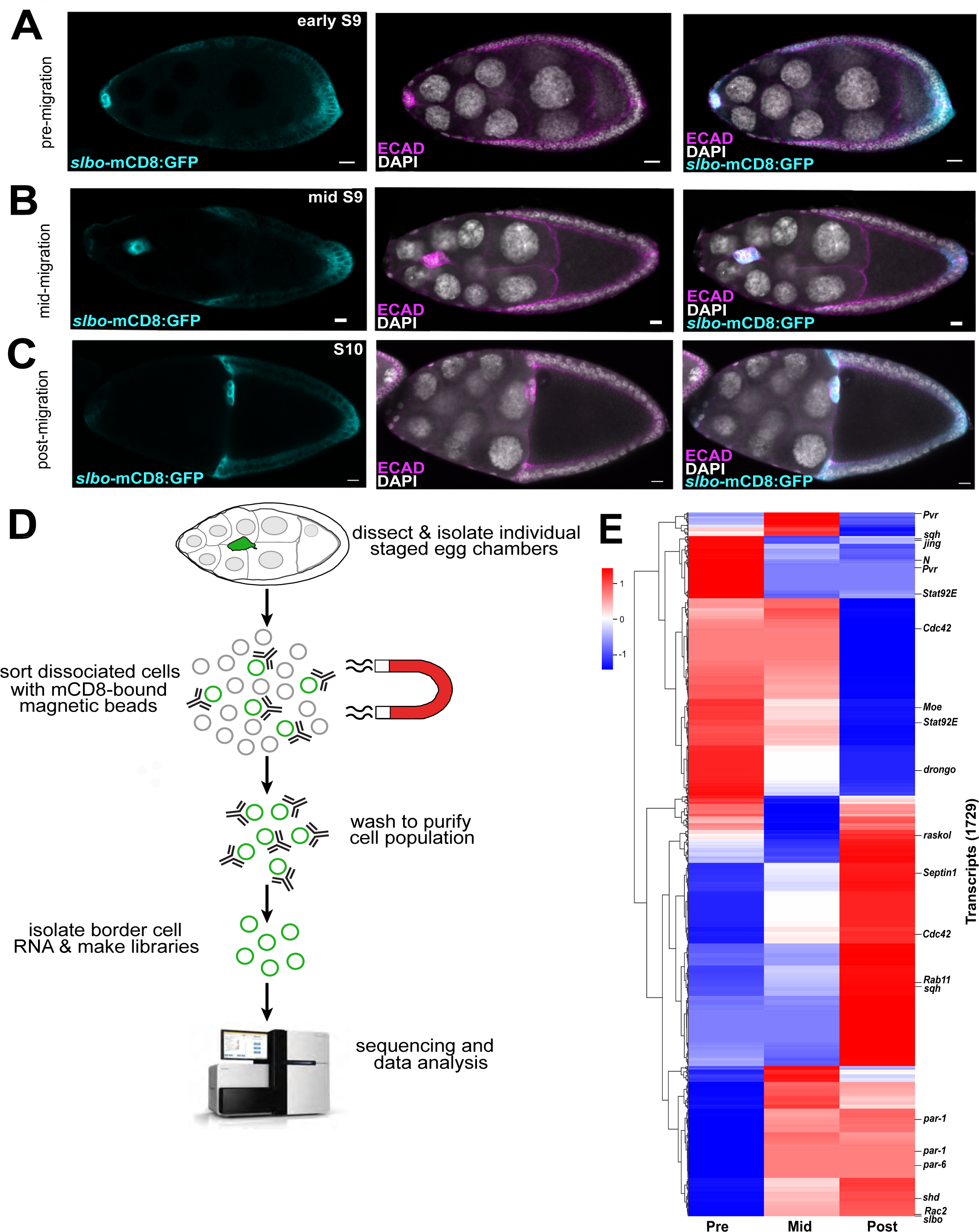
RNA sequencing of isolated populations of border cells reveals temporal changes in gene expression. (A-C) Representative egg chambers showing the patterns of *slbo-*mCD8:GFP expression in the border cell cluster prior to (A), during (B), and after migration (C). (D) Schematic of how the mCD8:GFP-positive border cells were sorted and selected using mCD8 magnetic beads. RNA from isolated border cells was then prepared for RNA sequencing and downstream analyses. (E) Heatmap of sequencing results, identifying significant differential expression for 1,729 transcripts from isolated border cells during their migration (EBSeq-HMM FDR<0.05). Several border cell migration-related genes are highlighted. Scale bars represent 10 μm. All significantly differentially expressed transcripts are shown in **Supplemental Table 3**. HiSeq 2500 graphic courtesy of Illumina, Inc.

The formation and migration of border cells requires multiple signals that coordinate dynamic cellular behaviors. First, border cells are specified and recruited by Janus kinase/signal transducer and activator of transcription (JAK/STAT) signaling [14; 15; 7; 16]. The anterior polar cells secrete a cytokine-like ligand, Unpaired (Upd1), which activates JAK/STAT in a gradient within the adjacent follicle cells. Cells with highest levels of JAK/STAT activity turn on a critical downstream transcription factor, the C/EBP ortholog Slow border cells (Slbo). Slbo activates a number of genes which together induce cells to become migratory border cells [17; 12; 7; 18]. A key target of Slbo is E-cadherin, which promotes the ability of border cells to migrate upon and between the nurse cells [19]. E-cadherin also keeps border cells adhered to each other and to the polar cells, particularly as the cluster navigates the dense and crowded tissue environment [20; 21; 22; 23]. Continuous JAK/STAT activation is required throughout migration to keep border cells motile, although how it does so is still unclear [24]. Second, a pulse of ecdysone at stage 9 of oogenesis activates the Ecdysone receptor (EcR) and the transcriptional co-activator Taiman (Tai) so that border cells begin their migration at the right time [25; 26; 27; 28]. Ecdysone helps localize E-cadherin within the cluster but also stimulates the expression of multiple genes that promote migration [25; 28]. Third, once specified, border cells are guided to the oocyte by secreted chemoattractant ligands from the oocyte that activate two receptor tyrosine kinases on border cells, PDGF- and VEGF-receptor related (PVR) and Epidermal Growth Factor Receptor (EGFR) [29; 30; 31; 32]. PVR and EGFR together stimulate the formation of dynamic F-actin-rich protrusions at the front of the cluster via the small GTPase Rac [30; 8; 33; 34]. Extension and retraction of these protrusions help the cluster navigate between the nurse cells and move posteriorly towards the oocyte [35; 22; 8]. Additional signaling pathways actively contribute to keeping border cells organized and motile. For example, Jun-kinase (JNK) helps border cells stay cohesive through regulation of the polarity protein Par-3 (Bazooka; Baz), whereas Hippo/Warts polarizes F-actin to the periphery of the cluster through the actin-regulatory proteins Enabled (Ena) and Capping protein [36; 37; 38].

Many of these signals and their downstream targets are conserved, making border cells a powerful model to identify additional genes that promote distinct aspects of collective cell migration. Previous microarray analyses of isolated border cells, along with multiple genetic screens, have uncovered genes expressed in and required for border cell formation and collective movement [17; 28; 5; 6; 18]. These expression screens pooled isolated border cells at all stages of the migration process, and thus were unable to determine if specific genes, or networks of genes, were dynamically expressed. Moreover, newer technologies have improved both the specificity and dynamic range of identifying transcripts. Therefore, we sought to comprehensively identify molecular determinants of dynamic border cell migration behaviors. In this study, we sequenced RNA isolated from border cells at three specific stages of migration: pre-migration, just prior to delamination; mid-migration, during movement of border cells between the nurse cells; and post-migration, when border cells reached the oocyte. We report here that 1,729 transcripts from 1,394 unique genes are temporally and differentially expressed at the three migration stages. These genes fall into nine clusters of co-expressed genes. Further gene ontology analyses and constructed protein-protein (PPI) interaction networks identify multiple categories of differentially expressed genes. In addition to genes expected to function in collective cell migration, such as those that regulate the cytoskeleton, cell adhesion, and tissue morphogenesis, we unexpectedly found an enrichment of genes involved in immune signaling, stress response pathways, and ribosome biogenesis. Finally, we characterized and confirmed the *in vivo* expression and functions of a subset of the identified genes in migrating border cells, thus validating this approach. Together, our results highlight new networks of genes that are differentially expressed during border cell migration and that represent potential regulators of dynamic collective cell behaviors.

## Results

### Transcriptional profile of temporally expressed genes at three border cell migration stages

Previous studies identified genes enriched in border cells using microarray analyses [17; 18]. However, whether some genes are differentially expressed at different stages of migration was unknown. Moreover, a subsequent analysis found that only 10% of genes between these two border cell microarray experiments overlapped [39]. Thus, our current picture of the dynamics of transcripts expressed in migrating border cells is incomplete. Therefore, we sought to identify temporally-expressed genes at three distinct stages of border cell migration: pre-migration, when border cells have rounded up into a cluster and have extended a protrusion in the direction of migration, but have not yet left the follicle cell epithelium (**Figure 1A**); mid-migration, when border cells have delaminated and migrated anywhere along the migration pathway (**Figure 1B**); and post-migration, when border cells have finished their migration at the oocyte boundary (**Figure 1C**). We performed bulk RNA sequencing of border cells isolated at these three distinct stages using a previously validated method of magnetic-bead cell sorting (*see* Methods; [18]). The *slbo* enhancer was used to directly drive mCD8:GFP in border cells (**Figure 1A-C**). A few additional follicle cells also express *slbo-*mCD8:GFP, including centripetal cells at stage 10 (**Figure 1C**). Egg chambers at the relevant stage were manually sorted using the GFP fluorescence, dissected, and pooled, followed by isolation and cell sorting of individual mCD8:GFP-expressing cells (**Figure 1D**). Three biological replicates were performed for each stage of migration, except the pre-migration stage in which one replicate failed due to low levels of RNA in the sample. RNA sequencing of these sorted border cell populations was then performed (**Figure 1D**).

After mapping the reads to the genome, expression for 25,645 of 30,504 *Drosophila* transcripts was detected in the genome (FlyBase version FB2021_02 Dmel Release 6.39; [40; 41]). As expected, GFP expression was observed from *slbo-*mCD8:GFP but did not significantly change in levels among samples, indicating consistency across the biological replicates (**Supplementary Table 1**). A total of 1,729 transcripts from 1,394 unique genes at the three different migration stages were determined to be differentially expressed with statistical significance (EBSeq-HMM FDR<0.05; **Figure 1E**; **Supplemental Tables 2 and 3**). As a positive control, expression of *slbo* (FBtr0072272), a gene known to regulate border cell identity and migration, was found to increase in expression during border cell migration (**Figure 1E, Supplemental Table 3**). A significant number of genes known to be expressed in border cells and required for their migration had differing levels of expression from pre- to post-migration (**Figure 1E**). Border cell-related genes that changed expression levels during migration included *drongo*, *Pvr*, *Rab11, Rac2, Raskol, Septin1*, and *spaghetti squash* (*sqh*) [42; 5; 23; 6; 43]. Additional genes such as *jing*, *Notch* (*N*), and *Stat92E*, known to be primarily expressed or activated in border cells at these stages, were also differentially expressed [44; 45; 24; 46]. These data support the idea that while other follicle cells were potentially isolated, relevant genes for border cell migration were enriched due to consistently higher levels of *slbo-*mCD8:GFP in border cells, and thus enrichment of border cells in the isolated cell populations (**Figure 1A-C**; *see* Methods).

We next asked if these temporally expressed genes overlapped with other genes known to be expressed in border cells or required for their migration. We compiled a comprehensive list of genes required for border cell migration from the published literature (**Supplemental Table 4**). Using the entire sequencing dataset, we then analyzed if any of these genes were differentially expressed. We identified 496 border cell migration transcripts that exhibited significant differential expression from pre- to post-migration stages (**Figure 2A; Supplemental Table 4**). We next compared our sequencing data to two previous microarray gene expression studies, which identified genes that were up- or down-regulated in border cells relative to other follicle cells and compared to *slbo* mutant border cells [17; 18]. Both microarray studies assessed the expression of genes in border cells that were isolated from whole ovaries but did not perform temporal staging of egg chambers. Thus, it was unknown if any genes were temporally expressed. Our analyses found that 1,145 transcripts from the Borghese et al. [17] study, representing 85.8% of total genes within the microarray, exhibited distinct patterns of expression during border cell migration (**Figure 2B; Supplemental Table 4**). Similarly, we found 853 transcripts, representing 71.7% of identified genes from the Wang et al. [18] microarray, that were differentially expressed (**Figure 2C; Supplemental Table 4**). Next, we asked whether genes that might be expected to function in border cell collective migration or development were temporally expressed from pre- to post-migration stages. Collectively migrating cells, including border cells, require the actin cytoskeleton and adhesion for movement and to organize the collective [5]. Indeed, 154 actin cytoskeleton transcripts and 147 adhesion transcripts were differentially expressed (**Figure 2D, E; Supplemental Table 4**). These included actin binding genes *Gelsolin (Gel)*, *Zasp52,* and *Vinculin (Vinc),* and adhesion-related genes *rhea, p120 catenin (p120ctn),* and *Fasciclin 1 (Fas1).* We also found 191 transcripts associated with epithelial-mesenchymal transitions (EMT) that exhibited differential expression during migration (**Figure 2F; Supplemental Table 4**). Even though border cells retain many epithelial features, including high levels of apical-basal cell polarity proteins and E-cadherin, and thus do not undergo true EMT [5], genes such as *Goosecoid (GSC), snail (sna),* and *twist (twi)* were differentially expressed in border cells [47; 48]. Transcripts of genes associated with oogenesis and transcription were also differentially expressed in migrating border cells (**Figure 2G, H; Supplemental Table 4**).

**Figure 2.**
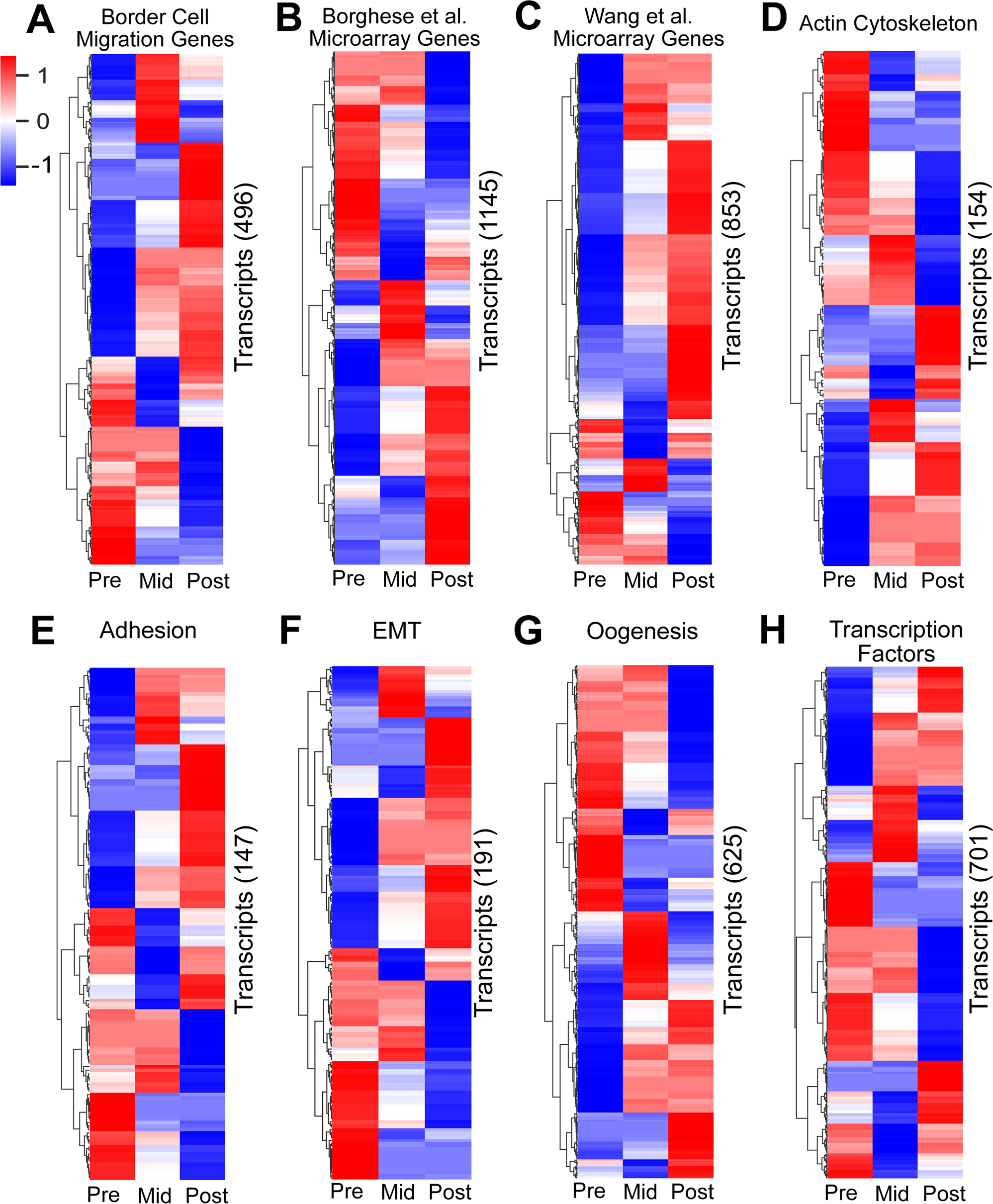
Migration and development-related genes are temporally expressed during border cell migration. (A-H) Heatmaps of significantly differentially expressed transcripts (EBSeq-HMM FDR<0.05), focused on subsets of migration-related (A-F) or developmentally-related (G, H) gene categories. The primary literature or other databases were used to identify genes and GO terms (*see* Methods for details). Heatmaps represent differential expression in border cells at pre-, mid- or post-migration. A *z*-score of 1 (shown in red) signifies up-regulation; a *z*-score of −1 (shown in blue) signifies down-regulation. (A-C) Transcripts for genes known to be required or expressed in border cells during their migration (A), or shown to be significantly upregulated in border cells versus non-migratory follicle cells from two microarray studies, [17] (B) and [18] (C). (D-H) Transcripts for migration-related (D-F) or developmentally-related (G, H) genes, including the actin cytoskeleton (D), adhesion (E), epithelial-to-mesenchymal transition (EMT; F), oogenesis (G), or transcription factors (H). All data are shown in **Supplemental Table 4**.

### Identification of distinct patterns and classes of co-expressed genes during border cell migration

To better understand the molecular control of border cell migration, we next used *clust* to determine which of the significantly differentially expressed genes were co-expressed from early to late border cell migration. *Clust* analysis automatically extracts optimal gene co-expression patterns in RNA-seq data that have a high correlation with similar biological activity [49]. Here, our *clust* analysis resulted in nine distinct clusters of temporally co-expressed genes (**Figure 3**). Of the transcripts with significant differential expression, 587 were grouped into three distinct co-expressed clusters with various patterns of increased expression across the migration stages (C0, C1, and C8; **Figure 3A, B, I**). These clusters included regulators of border cell migration such as *spaghetti squash* (*sqh*; C0), *big bang (bbg*, C1), and *slbo* (C8), (**Supplemental Table 5**). Another 556 transcripts were grouped into co-expressed clusters with various patterns of decreased expression across the migration stages (C4, C5, and C6; **Figure 3E, F, G**). Several genes known to be expressed or required in migrating border cells were found in these clusters, including *singed* (*sn*; C4)*, par-6* (C4)*, STAT92E* (C5), and rolling pebbles (*rols*; C5) (**Supplemental Table 5**). Finally, the remaining 119 transcripts sorted into three co-expressed clusters with variable expression patterns that specifically increased or decreased their expression only at mid-migration stages (C2, C3, and C7; **Figure 3C, D, H**). Known border cell migration regulators *Patj* (C2) and *Rac2* (C7) are found in these clusters, along with F-actin regulators (*HSPC300* in C3, *Arpc2* in C7) and several RhoGEFs (*RhoGEF2* and *RhoGEF3* in C3). These findings suggest that, despite their unique expression patterns and fewer transcript numbers, the three variable gene expression clusters are still biologically relevant (**Supplemental Table 5**).

**Figure 3.**
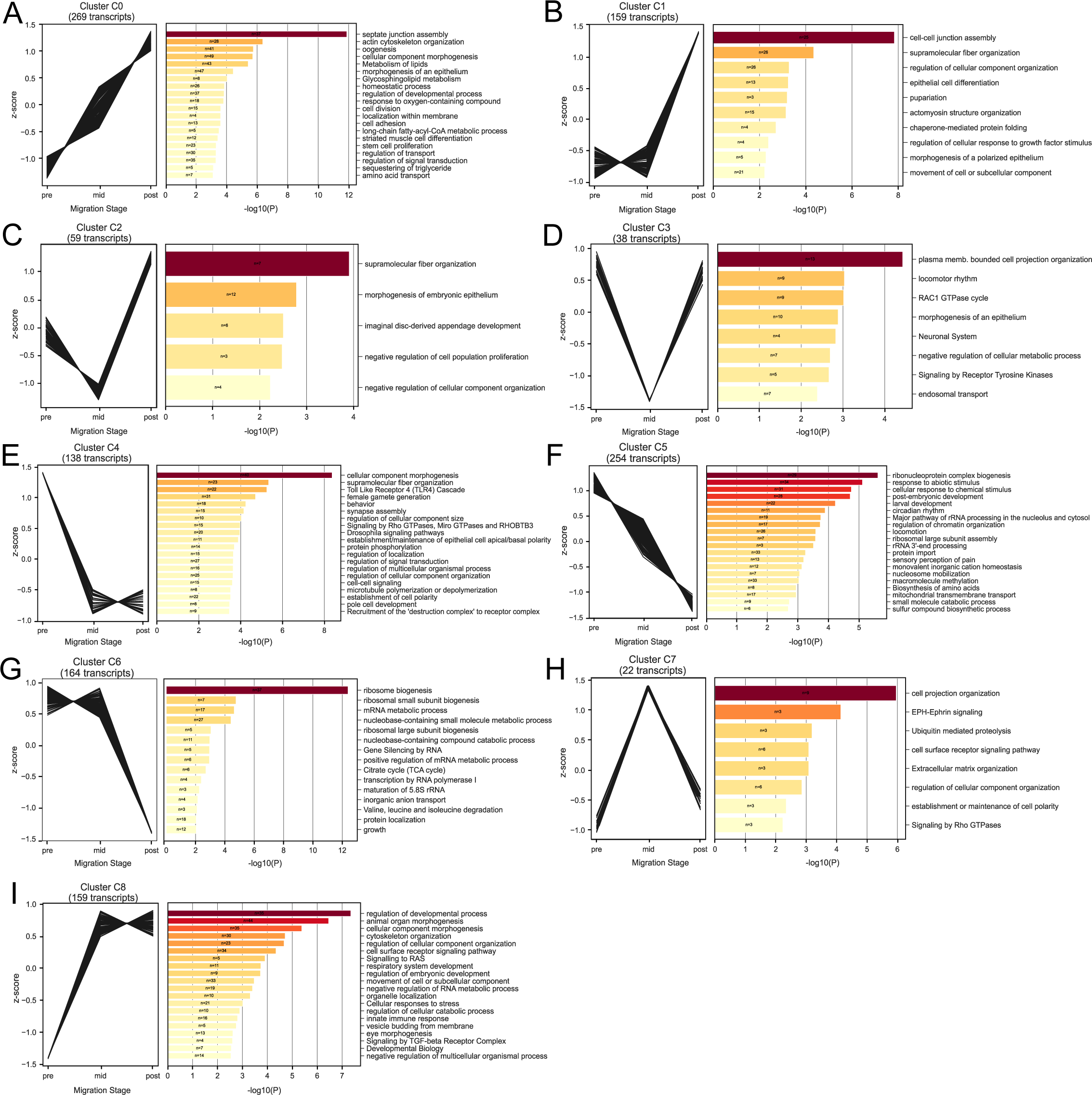
Differentially expressed transcripts are co-expressed during border cell migration and enriched for shared biological functions. (A-I, left graphs) Significantly differentially expressed transcripts sorted by *clust* into shared co-expression patterns in pre-, mid-, and post-migration stages. (A-I, right graphs) Metascape pathway and process enrichment analysis results for each co-expression cluster, showing the most significantly-enriched terms. *N,* number of genes enriched for a given annotation term; bars show significance of annotation terms, sorted by *p* values (-log10P; darker color, more significant values). Note that genes can be found in multiple Metascape annotation categories. Clusters C0 (A), C1 (B), and C8 (I) show patterns of increased expression during border cell migration. Clusters C4 (E), C5 (F), and C6 (G) share patterns of decreased expression during border cell migration. Clusters C2 (C), C3 (D), and C7 (H) share patterns of stage-specific increased or decreased expression at mid-migration stages. All data are shown in **Supplemental Tables 5 and 6**.

To obtain further insight into potential biological functions of the significantly differentially expressed clusters of genes, we performed a Metascape enrichment analysis [50]. This analysis identified top significantly enriched gene annotation terms, including those from gene ontology (GO), KEGG, Reactome, and other databases, for each clustered group of genes (**Figure 3A-I; Supplemental Table 6**). Clusters with increased expression from pre- to post-migration were notably enriched for genes that had four broadly shared functions (C0, C1, and C8; **Figure 3A, B, I**), specifically cell-cell junction regulation (C0: septate junction assembly; C1: cell-cell junction assembly), cytoskeletal organization (C0: actin cytoskeleton organization; C1: supramolecular fiber organization and actomyosin structure organization; C8: cytoskeleton organization), morphogenesis (C0: cellular component morphogenesis and morphogenesis of an epithelium; C1: regulation of cellular component organization and epithelial cell differentiation; C8: animal organ morphogenesis, cellular component morphogenesis, and regulation cellular component organization), and developmental processes (C0: oogenesis; C1: pupariation; C8: regulation of developmental processes). Clusters with decreased expression from pre- to post-migration were enriched for genes that shared a variety of functions ranging from development and signaling to basic cellular metabolic processes, such as ribosome biogenesis, rRNA processing, and ribonucleoprotein complex biogenesis (C4, C5, and C6; **Figure 3E-G**). The clusters with variable expression patterns at mid-migration were enriched for genes expected to function in cell migration (C2, C3, and C7; **Figure 3C, D, H**), such as regulation of morphogenesis (C2: morphogenesis of embryonic epithelium; C3: morphogenesis of an epithelium), regulation of cell projections (C3: plasma membrane bounded cell projection organization; C7: cell projection organization), and cytoskeletal organization (C2: supramolecular fiber organization; C3: Rac1 GTPase cycle). Together, these analyses highlight the complexity of border cell migration, with both expected (e.g. cytoskeletal genes, morphogenesis) and unexpected (e.g. metabolic processes) differentially co-expressed genes.

### Interaction networks reveal enrichment of migration-related, immune signaling, and ribosome biogenesis genes

The above-described *clust* analysis positioned the significantly co-expressed genes into clusters with predicted or known annotated functions, but did not capture relationships such as protein-protein interactions (PPI) among the gene products. Specifically, we wanted to understand how the proteins encoded by the differentially expressed genes might interact during border cell migration. Therefore, we examined potential PPI networks present in each of the *clust* co-expression clusters during border cell migration. We specifically focused on those co-expressed clusters with at least 150 significantly differentially expressed genes, namely clusters C0, C1, C4, C5, C6, and C8 (**Figure 3**). To identify PPI networks, we used Cytoscape to integrate Metascape annotations, which includes FlyBase PPIs, and STRING-based PPIs (**Figure 4; Supplemental Figure 1**; *see* Methods) [51; 52]. The PPI networks were further refined by manual curation using functional data from FlyBase to enhance the predictions of biological functions **(Supplemental Table 7)**. Demonstrating the utility of this approach, we found that at least 40% of the genes in each of these co-expressed clusters contained known physical protein interactions.

**Figure 4.**
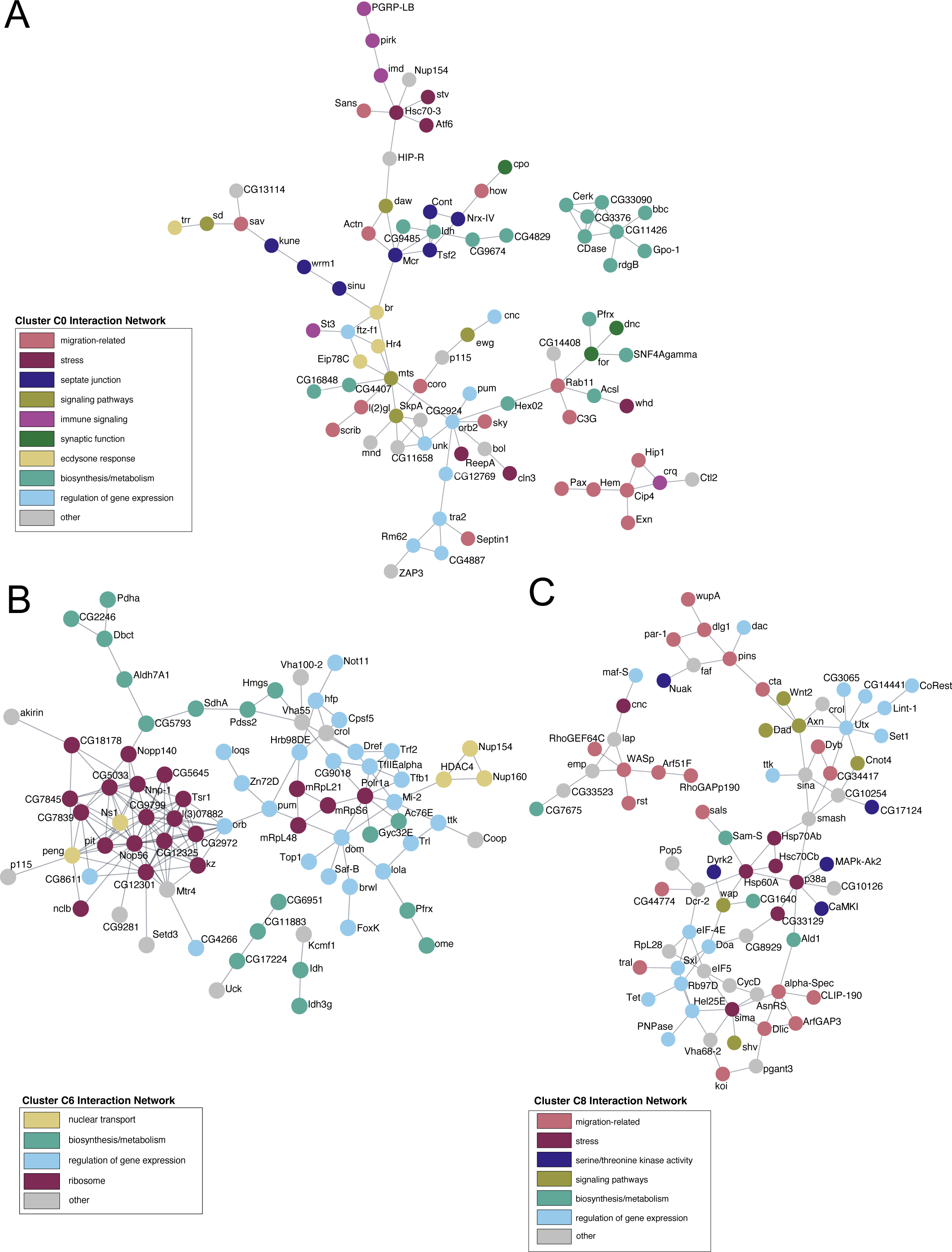
Differentially co-expressed genes form protein-protein interaction networks. Physical protein interaction (PPI) network analysis of gene products from selected co-expression clusters. Functional annotation keywords were used to assign color to proteins in the networks. (A) Co-expression cluster C0, encompassing genes that increase expression during migration, contains PPI nodes with migration-related functions (pink), including functions known to regulate border cell migration such as septate junction regulation (dark blue) and ecdysone response function (mustard). Individual networks consist of proteins with biosynthesis/metabolism (cyan, upper right) and migration-related (pink, lower right) functions. (B) Co-expression cluster C6, encompassing genes that decrease during migration, forms large nodes for ribosome function (magenta), biosynthesis/metabolism (cyan), and regulation of gene expression (light blue). Individual networks consist of proteins with biosynthesis/metabolism (cyan, lower center). (C) Co-expression cluster C8, encompassing genes that increase expression, forms one protein interaction network with migration-related (pink), multiple signaling pathways (light green), and gene expression regulation (light blue) functions. One individual network consists of proteins with migration-related and additional categories (upper left). For each PPI network, the “other” category (gray) either denotes genes that do not have available FlyBase annotations/data or for which three or fewer genes were annotated. All data, including annotations and keywords, are shown in **Supplemental Table 7**.

From this analysis, we identified major interaction networks within each of these significantly differentially expressed clusters (**Figure 4; Supplemental Figure 1; Supplemental Table 7**). We first analyzed networks of protein-protein interactions found in three clusters with patterns of increasing expression, C0, C1, and C8. Two of these co-expression clusters, C0 and C8, had modules of protein interactions with functions known or predicted to be important for collective cell migration (**Figure 4A, C**). These migration-related activities included regulation of the cytoskeleton, cell adhesion, small GTPase activity, and cell polarity. Cluster C0 had additional functional interaction modules with proteins required for septate junction formation and proteins involved in ecdysone signaling response, both of which are required for border cell migration (**Figure 4A**) [53; 25; 27; 54]. Cluster C8 was further enriched with modules of proteins with annotated broad functions in development, including cell signaling, serine-threonine kinase activities, and regulation of gene expression (**Figure 4C**). Notably, both C0 and C8 also included proteins with annotated functions in stress signaling (**Figure 4A and 4C**). The third co-expression cluster with a pattern of increasing expression, C1, had fewer protein interactions (**Supplemental Figure 1A**). However, the C1 nodes still formed functional modules with predicted or known migration-related activities, along with cellular transport and biosynthesis/metabolism. Next, we analyzed networks of protein interactions in the three co-expression clusters with various patterns of decreasing expression, C4, C5, and C6. Two of these co-expression clusters, C4 and C5, had modules of protein interactions with migration-related functions (**Supplemental Figure 1B and 1C**). Both C4 and C5 also had modules of proteins with functions that regulate gene expression and cell signaling. Cluster C6 was enriched for functional modules of proteins that regulate gene expression and proteins involved in biosynthesis and metabolism (**Figure 4B**). Overall, the assembled protein interaction networks reveal functions predicted or known to be important in border cell migration, but also new and unexpected functions.

The network analysis identified two major molecular pathways not typically associated with collective cell migration, immune response regulation in cluster C4 and ribosome-related functions in clusters C5 and C6 (**Figure 4B**; **Supplemental Figure 1B, C; Supplemental Table 7**). To enhance this analysis, we investigated if additional genes involved in immune response or ribosome-related activities were enriched throughout the co-expression networks. We used Metascape and GO enrichment categories (**Supplemental Table 6**) to curate genes with annotated “immune” or “ribosome” functions from each of the differentially co-expressed clusters (**Supplemental Tables 8 and 9**). We used additional pathway and function information from FlyBase and the primary literature to assemble these genes into graphical maps of immune signaling (**Figure 5A**; **Supplemental Figure 2**) and ribosome functions (**Figure 5B**).

**Figure 5.**
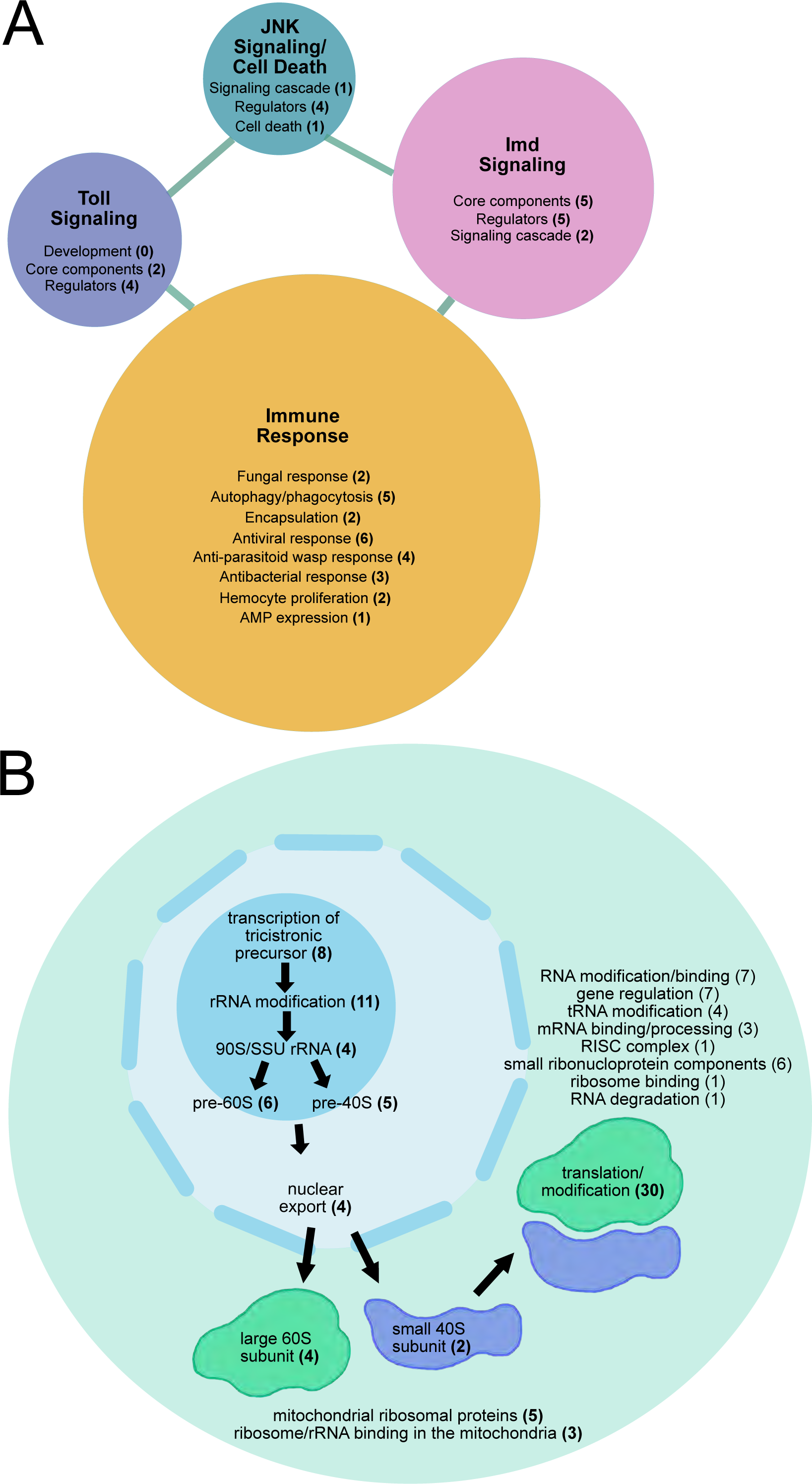
Networks of immune signaling and ribosome biogenesis genes differentially expressed during border cell migration. Graphical representation of the networks of differentially co-expressed genes in border cells annotated with immune (A) or ribosome-related (B) functions. (A) Co-expressed genes enriched for immune functions were sorted into Toll Signaling, Imd Signaling, JNK Signaling/Cell Death, or Immune Response categories. Genes involved in defense against fungus, virus, bacteria, or parasitoid wasp, as well as genes implicated more broadly in immune cell or cell death functions, but not linked to a signaling pathway, comprise the “immune response” category. (B) Co-expressed genes enriched for ribosome function were sorted into ribosome biogenesis in the nucleolus, nuclear export, large and small ribosomal subunit, translation/modification and other categories as indicated. For simplicity, the total number of genes enriched for small ribonuclear protein component functions is shown in the cytoplasm; a subset of these genes have predicted or demonstrated nuclear localization. All data are shown in **Supplemental Table 8** (A, immune functions) and **Supplemental Table 9** (B, ribosome functions).

Multiple immune pathway genes were identified. These genes were mainly found in co-expression clusters C4, C0, and C8, which had patterns of increasing or decreasing expression (**Figure 5A; Supplemental Figure 2; Supplemental Table 8**). The largest number of genes were classified as having general immune response functions. These 21 genes included those with specific functions in fungal, antiviral, or antibacterial responses, and those with less specific immune roles such as phagocytosis, encapsulation, or hemocyte proliferation (**Figure 5A**; **Supplemental Table 8**). Importantly, we identified core components and genetic regulators of both the *Drosophila* Toll and Immune Deficiency (Imd) innate immune signaling pathways (**Figure 5A; Supplemental Figure 2A, B**). Six genes involved in the Toll pathway and twelve genes involved in the Imd pathway were differentially expressed in border cells. Some critical genes in the canonical Toll cascade such as Toll itself, Cactus, and Dorsal, which also promote embryonic development, were not differentially expressed in border cells [55; 56]. The immune-specific transcription factor Dorsal-related immunity factor (Dif) was also not differentially expressed. However, other key genes such as the adaptor protein *Myd88* and the ligand *spatzle* (*spz*) in the Toll pathway and the adaptor protein *Imd* and the downstream transcription factor *Relish* (*Rel*) in the Imd pathway were differentially expressed in border cells (**Supplemental Figure 2A, B**). Jun-kinase (JNK) signaling contributes to the immune response by promoting differentiation of immune cells and sensing stresses such as infections, triggering production of antimicrobial peptides, and promoting wound healing [57]. JNK can also have other functions, including regulating polarity and cluster cohesion in border cells [37; 58]. Nonetheless, five regulators of the JNK signaling pathway were significantly differentially expressed in border cells (**Figure 5A; Supplemental Figure 2C**). Thus, some, though not all key members of the innate immune pathways, along with known regulators, are differentially expressed during border cell migration.

A substantial number of genes with ribosome functions were identified (87 in total). Ribosomal-related genes were primarily found in co-expression clusters C5 and C6, with decreasing patterns of expression during migration, with only four genes exhibiting increased expression (**Figure 4B; Supplemental Figure 1C; Supplemental Table 9**). Ribosome biogenesis primarily takes place in the nucleolus and requires hundreds of proteins to assemble the ribosomal subunits [59]. The greatest number of genes found in border cells function in ribosome biogenesis within the nucleolus (**Figure 5B; Supplemental Table 9**). These genes (34 in total) encode proteins annotated to regulate transcription of ribosomal RNA (rRNA), modify rRNA, or form the immature large and small ribosomal subunits. Another four genes encode proteins that export ribosomes from the nucleus to the cytoplasm. Genes with annotated functions in the maturation of the ribosomal subunits after export to the cytoplasm (six genes) or mitochondrial ribosome function (eight genes) were also differentially expressed. Additional differentially expressed genes include those that encode the large and small ribosomal subunits, function in the mature ribosome to regulate cytoplasmic translation/modification, and methylate/bind RNA, as well as other various functions (**Figure 5B; Supplemental Table 9**). These data together indicate that ribosome biogenesis genes are differentially expressed and broadly decrease from early to late border cell migration.

### Genes identified by transcriptomics are expressed in migrating border cells *in vivo*

The transcriptomic analyses described above revealed large sets of significantly differentially expressed genes. We wanted to further validate the expression of a subset of these genes in migrating border cells *in vivo*. To do this, we took advantage of an open repository of ovarian RNA fluorescence *in situ* hybridization images, the Dresden Ovary Table [60; 61]. Of the 1,262 significantly differentially expressed genes (**Figure 3**), 17% had available DOT FISH images at oogenesis stages 8-10 when border cells form and migrate (**Supplemental Table 10**). To validate this approach, we selected two genes known to be expressed in border cells [17; 12; 18], *slbo* (C8) and *sn* (C4), that were also found to be significantly differentially expressed in the *clust* datasets. As expected, both *slbo* and *sn* transcripts were enriched in border cells at pre-, mid-, and post-migratory stages (**Figure 6A-B, M**; **Supplemental Table 10**). Two additional genes with known functions in border cells, *rols* and tramtrack (ttk), also had strong expression in border cells at these stages (**Supplemental Table 10**) [17; 18]. Fourteen additional genes from various enriched clusters showed strong evidence of border cell expression from stages 8 to 10 (**Figure 6M**; **Supplemental Table 10**), including *CG11147* (**Figure 6C**), *couch potato* (*cpo*; **Figure 6D***), Neprilysin 2* (*Nep2*; **Figure 6E**), and *Cadherin 74A* (*Cad74A*; **Figure 6F**). A further six genes had visible expression in border cells during at least one stage of migration: *B4* (**Figure 6G**), *fng* (**Figure 6H**), *CG11007* (**Figure 6I**), *Lipase 4* (*Lip4*; **Figure 6J***), CG17124* (**Figure 6K**), and *varicose* (*vari*; **Figure 6L**). It is possible that these genes are expressed at other stages of border cell migration but clear images for these genes in the DOT were limited. Moreover, the dynamic range of fluorescent RNA *in situ* hybridizations does not allow assessment of differing transcript levels from early to late migration stages. Nonetheless, these data provide visual confirmation that genes identified in our RNA sequencing datasets are expressed in migrating border cells *in vivo*.

**Figure 6.**
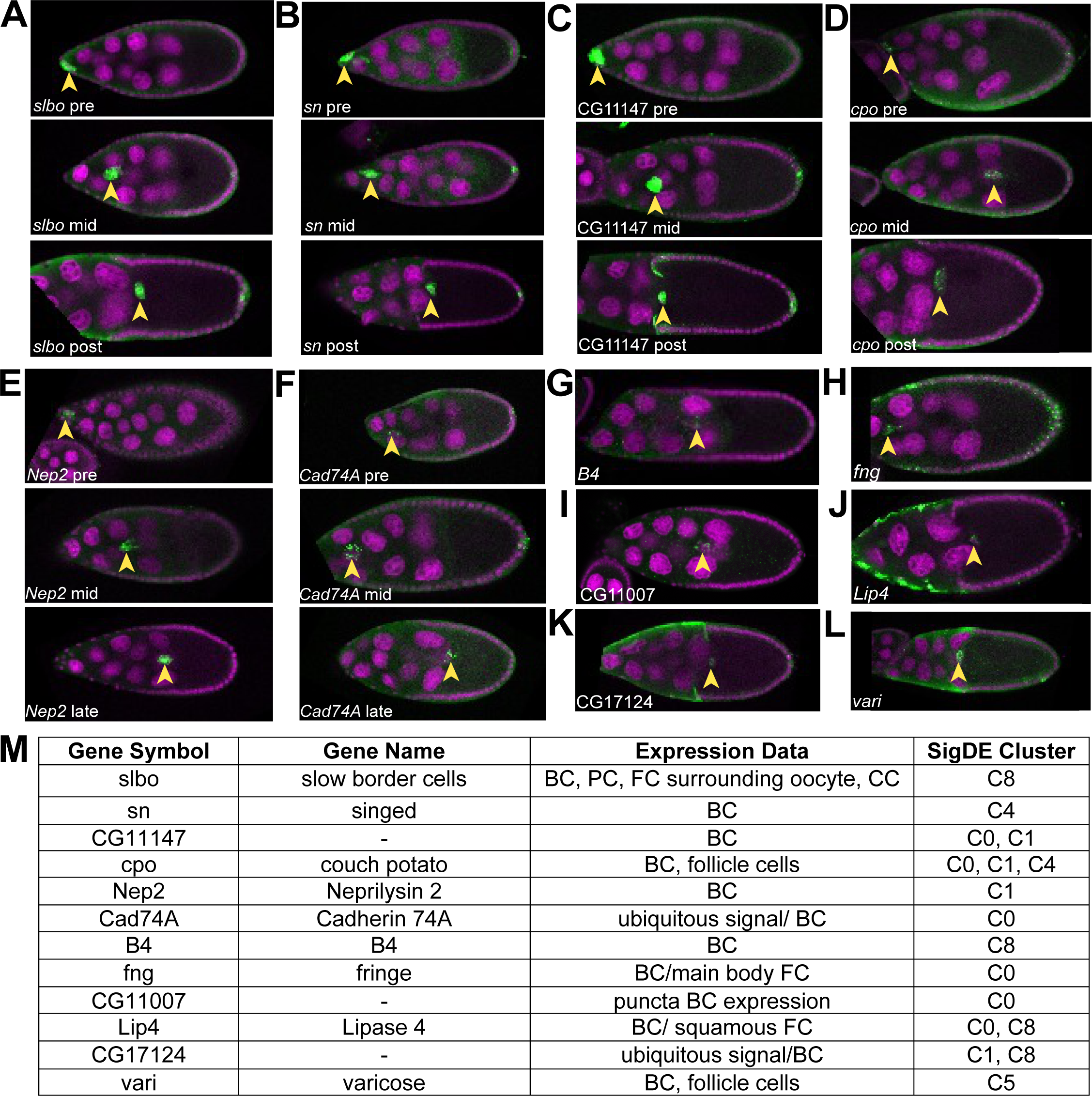
Expression patterns of genes in migrating border cells *in vivo*. Fluorescence RNA in situ hybridization patterns of genes from the transcriptomics analysis in migrating border cells using data from the Dresden Ovary Table (DOT) (http://tomancak-srv1.mpi-cbg.de/DOT/main). Representative images of stage 9 and/or stage 10 egg chambers were chosen for given genes of interest. RNA signal is in green and DAPI labels the nuclei in magenta. Anterior is to the left in all images. (A-F) RNA signal (green) is shown in pre-, mid-, and end (or late) migratory border cell clusters. (G-L) RNA signal (green) is shown at one stage of migration. (A, B) Expression of *slbo* (A) and *sn* (B), two genes known to regulate border cell migration. (C-L) Expression of multiple genes identified from the transcriptome analyses with previously uncharacterized roles in border cell migration. (M) Table of genes found in the DOT, along with their annotated expression patterns in the ovary, based on all available images, and location within one of the significantly differentially co-expressed gene clusters (SigDE Cluster). Arrowheads mark the position of border cells. All data are shown in **Supplemental Table 10**.

### Functional validation of temporally expressed genes in border cell migration

Finally, we wanted to functionally validate that temporally expressed genes in border cells were also required for their migration. We identified a subset of genes to be tested from the entire set of significantly differentially expressed clustered genes. Specifically, we tested genes with various GO terms that ranged from transcription, cell differentiation, adhesion, actin binding, and other various molecular functions, that were found across the different gene expression clusters (**Figure 3**; **Supplemental Table 11**). A total of 42 genes were knocked down by RNAi in border cells. We drove RNAi expression using the *c306-*GAL4 driver, which is expressed in border cells as well as in some anterior and posterior follicle cells (**Figure 7A-C**). Where possible, we tested at least two RNAi lines per gene. Migration was scored at stage 10 of oogenesis, by which time border cells have normally completed their migration at the oocyte anterior boundary (**Figure 7C-F**). Border cells that did not reach at least 76% of the distance to the oocyte were considered to have a migration defect (**Figure 7D, E**). As positive controls for this screen, we performed RNAi knockdown of two genes known to be required for border cell migration, *bazooka (Baz)* and *Rap1*, both of which impaired migration in ∼25% of egg chambers (**Figure 7D, E, G**) [62; 63; 64]. RNAi against *mCherry*, which encodes a fluorescent protein not normally found in *Drosophila*, was used as a negative control; *mCherry* RNAi did not significantly disrupt border cell migration (5% migration defects; **Figure 7E, F**). RNAi knockdown of 21 total genes (22 RNAi lines) resulted in border cell migration defects in >10% of egg chambers (**Supplemental Table 11**). Knockdown of eight of these genes resulted in stronger migration defects, with a range of 14-to-45% of border cells failing to complete their migration by stage 10 (**Figure 7E, H-K**). The genes identified here have a variety of predicted or known functions, including transcriptional regulation (*held out wings, how*; *Ches-1-like*; **Figure 7E**), small GTPase activity (*Arf51F*, also known as *Arf6*; **Figure 7E, H**), cell polarity (*serrano*, *sano*; **Figure 7E, I**), cell adhesion (*Basigin*, **Figure 7E, J**; *mspo*, **Figure 7E**), cell membrane organization (*Cip4*; **Figure 7E, K**), and other various functions (e.g., cell signaling, F-actin regulation, etc.; **Supplemental Table 11**). Knockdown of the transcription factor *Ches-1-like* had the strongest migration defects (**Figure 7E**; **Supplemental Table 11**). This RNAi line also has a predicted off-target match to *taiman* (*tai*). Tai is an Ecdysone Receptor coactivator that is required for border cell migration [25; 27; 28]. Thus, the migration defects seen by *Ches-1-like* RNAi could be due to knockdown of *tai* or reflect co-knockdown of both genes. However, a previous RNAi screen used an independent RNAi line to *Ches-1-like* with no predicted off-target matches and found that *Ches-1-like* knockdown completely prevented border cell migration [65], suggesting that *Ches-1-like* is required. The gene *karst*, which encodes βH-spectrin, was previously shown to help organize the border cell cluster [66]. Here we observed mild migration defects when *karst* was knocked down using either of two RNAi lines (**Figure 7E; Supplemental Table 11**). Knockdown of most genes resulted in mild migration defects, which could reflect incomplete knockdown due to RNAi efficiency or redundancy with other genes. Nevertheless, these data together demonstrate that at least a subset of the temporally expressed genes identified through RNA sequencing are required for normal border cell migration. Other genes identified here likely also function in border cells, but will require further work to determine their exact contributions.

**Figure 7.**
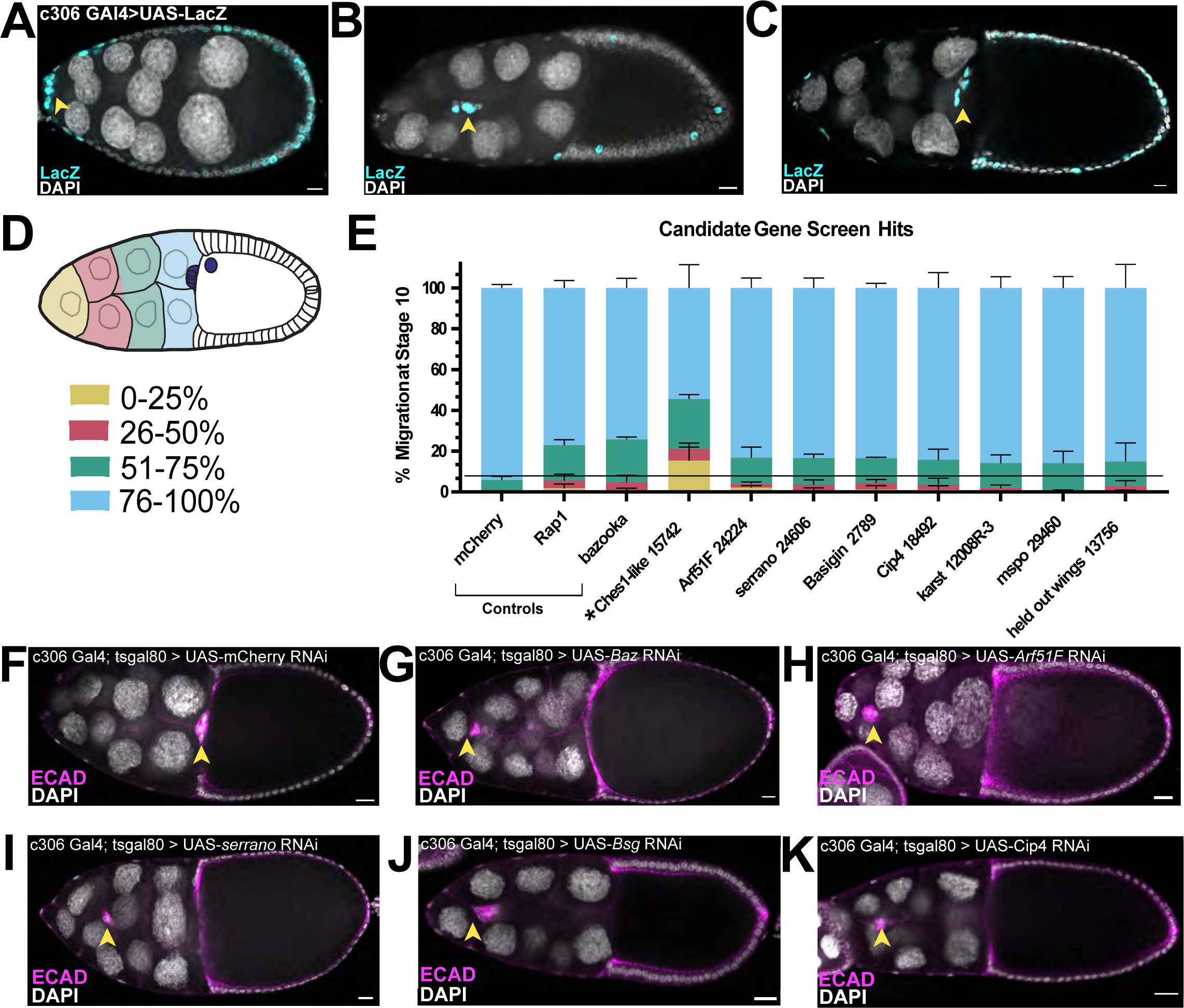
Functional assessment of temporally expressed genes in border cell migration. Validation of differentially expressed genes in border cell migration using RNAi. (A-C) Pattern of *c306*-GAL4, driving UAS-LacZ (cyan), in stage 9 pre-, mid-, and end migratory border cells and a few additional follicle cells. DAPI (gray) labels all nuclei in the egg chambers. (D-K) Knockdown of candidate genes driven by *c306*-Gal4; genotypes are *c306*-Gal4; tsGAL80 > UAS-RNAi. (D) Migration of border cells was scored based on the percentage of border cells that completed their migration (76-100%, blue) by stage 10 of oogenesis, by which time migration should be complete, or if the border cells stall along the migration pathway shown as the distance migrated away from the anterior tip of the egg chamber (0-25%, yellow; 25-50%, pink; 51-75%, green) as shown in the schematic. (E) Quantification of migration defects for genes from the transcriptome analyses, along with the negative control (*mCherry* RNAi) and positive controls *bazooka* (*Baz*) and *Rap1* RNAi, two genes known to regulate border cell migration. Shown are results for knockdown of 8 genes that resulted in significant migration defects at stage 10 (line); *Ches-1-like* (asterisk) RNAi had a strong migration defect but may partly be due to off-target effects. (F-K) Images of border cell migration at stage 10 for control (F, G) and experimental RNAi (H-K) egg chambers. Border cells (arrowheads) are labeled with E-cadherin (magenta); DAPI (gray) labels the nuclei of all cells. (F) Complete migration in a stage 10 *mCherry* RNAi control egg chamber. (G) RNAi knockdown of a positive control, *Baz,* to show representative migration defects. (H-K) Migration defects in *Arf51F* (H), *serrano* (*sano*; I), *Bsg* (J), and *Cip4* (K). Anterior is to the left in all images. Arrowheads mark the position of border cells. All scale bars represent 10 μm. All data and reagents are shown in **Supplemental Table 11**.

## Discussion

Border cell migration is a tightly regulated process in which multiple signaling pathways converge to specify cell fate and control dynamic cellular behaviors. This is reflected in our results from sequencing of transcripts from temporally staged populations of border cells. Here, we identified nine clusters of significantly differentially expressed transcripts with shared patterns of co-expression from pre- to post-migration. The Metascape and network analyses further revealed distinct molecular pathways and PPI networks that occur during various stages of the migration process. Our study found differential expression of genes involved in many cellular processes known or predicted to be important for collective migration. This included genes that function in the regulation of cell adhesions, cell and tissue morphogenesis, organization of the cytoskeleton, and cell polarity. In addition to these expected migration-related genes, genes that function in immune signaling, ribosome biogenesis, and stress response were also found to be differentially expressed in border cells. The relevance of a subset of differentially expressed genes was confirmed through RNA *in situ* patterns and by functional RNAi in border cells. Importantly, many of the differentially-expressed genes identified here overlapped with two previous microarray studies [18; 17]. However, these previous studies pooled all migration stages together and focused only on genes upregulated in border cells versus non-migratory follicle cells. The genes identified in our study thus represent a rich resource of gene expression networks that function during different stages of border cell migration and may inform our understanding of other cell collectives during development and in disease.

Among the top categories of differentially expressed genes in border cells were adhesion- and morphogenesis-related genes. A variety of adhesion proteins help collectively migrating cells stay together, maintain their cell shapes, communicate among cells, and move upon other cells and extracellular matrices [67; 68; 69]. Similarly, border cells require complex regulation of cell-cell adhesions for their collective migration to maintain cluster integrity and shape as well as to migrate on and between nurse cells [20; 67; 19; 23; 6]. Recent genetic screens and targeted approaches have uncovered various adhesion genes, including septate junction genes, that promote border cell collective cohesion and migration [53; 70; 23]. Consistent with this, we observed a broad pattern of dynamic adhesion-related gene expression from early- to late-migration stages. Septate junction assembly, cell-cell junction assembly, and cellular and epithelial morphogenesis genes were specifically enriched in co-expressed clusters whose expression increased from early to late migration stages. We further confirmed the expression and/or functions of several cell junction and morphogenesis genes, including *Cad74A*, *B4*, *fng*, *Arf51f*, *sano*, and *Cip4*. The adherens junction protein E-cadherin plays a major role in border cell cohesion and migration upon nurse cells [19; 20; 21], but surprisingly was not differentially expressed. However, we observed differential expression of *Stat92E* and *slbo*, which upregulate expression of *E-cadherin* just prior to border cell migration [19; 7]. Moreover, other regulators of E-cadherin protein localization and/or levels in border cells, such as *raskol*, *Hrb98de*, and *par-1* were also differentially expressed (**Figure 3**; [23; 71]). The identified differentially expressed cell adhesion, cell junction and morphogenesis genes are thus candidates to regulate critical features of border cell adhesion, cluster integrity, and migration.

We also identified differentially expressed “migration-related” genes within most of the co-expressed clusters and predicted PPI networks. These migration-related genes included those that encode regulators of the actin cytoskeleton, actomyosin structure, and regulators of Rho and Rac small GTPases. Border cells, like all migrating collectives, require finely-tuned Rac-dependent F-actin to extend and retract protrusions specifically at the cluster front for efficient movement [4; 2]. Non-muscle myosin II (myosin) contracts the rear to facilitate delamination from the epithelium [72; 43]. Actomyosin contraction via RhoA activation also helps border cells maintain an optimal cluster morphology as the group moves through the tight spaces between nurse cells [42; 73]. Regulators of actin and actomyosin were among the top annotated terms in the co-expressed clusters with increasing expression from early to late migration (e.g., clusters C0 and C1), consistent with their central roles in active cell migration. The differential expression of F-actin and myosin regulatory genes could help border cells quickly adapt to changing conditions in the immediate environment as border cells delaminate and move through the tissue. Supporting this idea, border cells actively respond to physical and chemical cues in the egg chamber by altering protrusion numbers, the shape of the collective itself, and the speed of their migration [74; 42; 73; 22; 34; 75; 8].

Genes important for cell polarity, along with genes associated with EMT, were also differentially expressed. Border cells do not undergo the stereotypical EMTs found in other migratory cell types during development or in cancer. Instead, border cells begin as epithelial cells with notable apical-basal polarity that is retained upon formation of the cluster and subsequent migration [63; 71; 76]. Border cells also upregulate E-cadherin, which is in contrast to cells that undergo complete EMT and downregulate E-cadherin [47]. Therefore, it was surprising that genes associated with EMT, including EMT-associated transcription factors *sna* and *twi*, were differentially expressed in border cells. Not all EMT genes play exclusive roles in EMT, however, and could have additional roles in cell migration. For example, two of the differentially expressed EMT genes, *crb* and *Notch*, also promote the polarity and cohesion of border cells [77; 46]. Moreover, EMT itself is quite complex [78; 48]. Many cells that undergo EMT during development and in cancer exhibit “plasticity” during their migration, and can transition back and forth between different epithelial and mesenchymal states [47; 48]. The high levels of apical-basal proteins in border cells promote various aspects of their migration, from delamination from the epithelium, extension of protrusions, follower cell behaviors, and maintenance of junctions between migrating border cells [63; 37; 71; 77; 76]. Supporting this role, we found that apical complex genes (e.g., *par-6*) and basal complex genes (e.g., *par-1*, *l(2)gl*, *dlg1*, and *scrib*) were differentially co-expressed and found in multiple PPI networks. Thus, while border cells retain epithelial polarity and require polarity genes for their migration, they may share features in common with other cells that undergo EMT.

Ribosome biogenesis and function genes were among the most highly represented genes whose expression was high early in border cells but decreased during the course of migration. The differentially expressed ribosomal genes included those that function in the transcription, processing, and modification of rRNAs and in the production and nuclear export of the small and large ribosomal subunits. Ribosome biogenesis is tightly linked to translation efficiency, which in turn regulates the levels of proteins. This suggests that border cells need high levels of proteins early in the migration process. In development, homeostatic levels of ribosomal biogenesis proteins and subsequent protein translation are critical, particularly for the specification of stem cells and germ cells and during times of tissue growth [79; 80; 81]. *Drosophila* germline stem cells require an increase in ribosome assembly, including higher expression of ribosome biogenesis and RNA processing genes, for proper growth and division [82; 80]. Conversely, altered ribosome biogenesis leads to defects such as skeletal and craniofacial abnormalities and to diseases termed “ribosomopathies” [79; 83, 80]. Such defects are due to some mRNAs being more sensitive than others to the overall availability of ribosomes [79].

Why might border cells need higher levels of ribosomal biogenesis genes during early phases of migration? Recent work indicates that increased levels of rRNA and ribosomes are important for migrating and invading cells. For instance, differential levels of ribosome biogenesis genes occur during the early phases of anchor cell invasion in *C. elegans* vulval development [84]. In this case, a burst in ribosome production coincides with an increase in the levels of pro-invasive proteins, which are needed for the anchor cell to breach the basement membrane [84; 85]. Similarly, in neural crest cells and breast cancer cells, rRNA and ribosome biogenesis is high and required to initiate the EMT program [86]. In some migrating cells, cellular protrusions such as lamellipodia have an enrichment of localized mRNAs, eukaryotic initiation factors, and ribosomes required for movement [87; 88; 89; 90]. Border cells similarly produce dynamic lead cell protrusions that detect guidance cues and help the cluster move between the nurse cells [22; 20; 34; 91]. As of yet, however, there is no evidence for localized enrichment of mRNAs or ribosomes in border cell protrusions. Nevertheless, an increase in ribosome biogenesis could support high levels of protein translation in border cells that leads to more efficient protrusion formation, delamination, motility, and/or collective invasion of the tissue. Interestingly, we found that the expression levels of ribosomal biogenesis genes significantly decreased by the end of border cell migration. Building the ribosome is energy intensive [92; 93; 59]. Thus, when border cells finish their migration, they may need to conserve energy and/or decrease their protein production. Further work will be needed to formally test how differing levels of ribosomal biogenesis genes contribute to border cell migration.

The transcriptome and PPI network analyses further revealed enrichment of genes whose protein products are annotated to function in immune signaling, in biosynthesis and metabolism, and stress response. Immune and biosynthesis/metabolic genes were found throughout all the major co-expressed clusters and networks. Thus, while differentially expressed, these genes were not associated with one specific stage of migration. Border cells express some, though not all, key components and regulators of the Toll and Imd signaling pathways, including genes that function in defense against viral, bacterial, fungal, or parasitic threats [56]. For example, Spatzle, Myd88, Imd and Relish were significantly differentially expressed during border cell migration, but Toll, Cactus, and Dorsal were not. The significance of these immune signaling genes is unknown since border cells are migratory epithelial cells and as such are not expected to have immune signaling functions. Interestingly, a previous RNAi screen found a requirement for multiple members of the Toll signaling pathway including Dorsal and Dif in border cell migration [65]. The biosynthesis/metabolic genes found to be differentially expressed in border cells have a wide variety of cellular functions. These functions include lipid metabolism and synthesis (e.g. Agpat3, SNF4Agamma), membrane homeostasis (e.g. CDase), and metabolic processes in mitochondria (e.g. Idh, Idh3g, SdhA). Stress response genes were found in networks within C0 and C8, two clusters that had increased expression from pre- to late-migration. Differentially expressed stress genes include those whose protein products regulate oxidative stress (e.g., Cnc; Whd) and the unfolded protein response (e.g., Atf6, Hsc70-3). Recent work has demonstrated roles for metabolic and mitochondrial genes in immune cell (hemocyte) invasion during *Drosophila* embryogenesis [94]. While intriguing, it remains to be determined whether the immune, biosynthesis/metabolism, and stress response genes identified here have specific roles in border cell development or migration, or if they mainly provide basal homeostatic functions.

## Conclusions

Collective cell migration is a highly dynamic process that requires intricate coordination of multiple developmental processes ranging from differential adhesion that keeps cells together, to a dynamic actin cytoskeleton that directs cell movements and migratory protrusions. Because of this complexity, it has been a challenge to understand the dynamic changes that occur in the molecular architecture of cell collectives as they migrate. Our transcriptome analyses, and functional validations, in border cells identified known biological processes in collective cell migration such as adhesion and polarity, but also implicated a novel role for the ribosome in border cell migration. Given the striking similarities between border cells and other cell collectives including cancer cells [95; 3; 96], the genes identified here represent a wealth of new candidates to investigate the molecular nature of collective cell migration.

## Methods

### Drosophila genetics

All fly stocks and crosses were maintained at 25°C, unless otherwise indicated. A *slbo-*mCD8:GFP/CyO stock (gift of X. Wang, CNRS/University of Toulouse, Flybase protein ID FBtp0002652) was outcrossed to *w^1118^* and used to obtain sorted border cell populations. For RNAi screening, stocks were obtained from the Bloomington Drosophila Stock Center (BDSC), the Vienna Drosophila Resource Center (VDRC), and NIG-Fly. Where possible, two independent non-overlapping RNAi lines were chosen per gene. Females from a *c306* Gal4; tsGAL80/CyO stock were crossed to UAS-RNAi males to drive expression of the knockdown construct in border cells. RNAi against *mCherry* (BDSC 35785) was used as a negative control, and *bazooka* (*Baz*; VDRC 2914) or *Rap1* (BDSC 57851) were used as positive controls. Female progeny from these crosses were selected and fattened on wet yeast paste overnight at 29°C to allow maximal GAL4/UAS expression and inactivation of tsGAL80. All RNAi lines are shown in **Supplemental Table 11**.

### Isolation of staged border cell RNA, library preparation, and Illumina sequencing

To isolate staged egg chambers for cell sorting, whole ovaries were first dissected from fattened 3- to 5-day old *slbo-*mCD8:GFP/+ females followed by dissection into ovarioles in room temperature “live imaging media” [Schneider’s media, pH 6.95 (Thermo Fisher Scientific),15% fetal bovine serum [Seradigm FBS; FBS (VWR), 200 µg/mL insulin (bovine pancreatic, Sigma-Millipore #I5500)] [97]. Stages 9 and 10 egg chambers [13] were manually selected in a two-well concave slide by observing GFP in the border cells (*slbo-*mCD8:GFP) on a fluorescent stereomicroscope. Egg chambers were considered to be at the “pre-migration” stage if border cells had rounded up and had a visible protrusion. Egg chambers with border cells that detached from the epithelium and had moved into the egg chamber anywhere along the path of migration, but had not reached the oocyte yet were considered “mid-migration”. Egg chambers with border cells that had reached the oocyte were considered “post-migration”. The stalk between egg chambers was cut and 40-60 GFP-positive egg chambers at each stage (pre-, mid-, and post-migration) were individually pooled and transferred to separate microcentrifuge tubes. Egg chambers were washed twice with cell dissociation buffer (Sigma-Millipore, cat. no. C5914-100ML), then treated with 10 mg/mL elastase (Sigma-Millipore SIGMA, cat. no. E0258-5MG; lot number SLBL4608V) in cell dissociation buffer for 30 min at 25°C with occasional agitation, followed by pipetting the mixture. Full cell dissociation was confirmed by visual inspection on a fluorescence stereomicroscope. The elastase reaction was stopped by addition of Schneider’s media. Cells were collected by centrifugation at 1000 rpm for 2 minutes and resuspended in PBS, 0.5 mg/mL Bovine Serum Albumin (BSA). Cells were washed twice in PBS supplemented with 0.5 mg/mL PBS prior to selection for GFP-positive cells.

Dissociated cells were incubated with anti-mCD8 antibody (Thermo Fisher Scientific, catalog number MA1-145) and 5 µl of protein-G Dynabeads Magnetic Beads (Thermo Fisher Scientific, #10003D) for 30 minutes at room temperature in the dark. Cells bound to the beads were isolated using a magnetic rack (Thermo Fisher Scientific, cat. no. 12321D) for 5 min at 25°C, followed by two washes with PBS supplemented with 0.5 mg/mL BSA. Magnetic isolation yielded ∼50-120 GFP-positive cells. The yield was assessed by fluorescence microscopy to ensure that all cells were GFP-positive. Captured cells were then suspended in 100 µl of Trizol™ (Thermo Fisher Scientific, cat. no. 15596026) and frozen at −20°C. RNA was isolated using the manufacturer’s instructions. The resultant purified RNA was stored in 75% ethanol at −80°C prior to library preparation. Each cell isolation yielded ∼50-100 ng of total high-quality RNA, as assessed by TapeStation analysis (Agilent Technologies, RIN*^e^* >8.0).

RNA-sequencing libraries were prepared with the Illumina TruSeq® version 2 stranded library preparation kit with 12 multiplexing barcodes. Prior to combination and loading, libraries were quantified by qPCR using a library quantification kit (NEBNext® Library Quantification Kit, E7630, New England Biolabs). Illumina sequencing was performed on a HiSeq2500 instrument for 100 single end read cycles (Genome Sequencing Core, University of Kansas).

### Read mapping, quantification, and normalization

Read mapping and quantification were performed using RSEM version 1.3.3 [98] into a Snakemake version 5.5.4 [99] pipeline. Illumina reads were trimmed for quality and adapter presence with Sickle version 1.33 Joshi2011-yt and Scythe version 0.994 Buffalo2011-za. Reads with quality less than PHRED 30 or less than 4 base pairs in length were discarded prior to mapping. Reads were checked for quality with FastQC version 0.11.6 Andrews2019-hi before and after trimming. Reads were mapped to the *Drosophila melanogaster* FlyBase predicted transcriptome (FB2021_02 Dmel Release 6.39; [41]), including splice variants, using bowtie RSEM included in the RSEM distribution Langmead2012-uk (for alignment statistics, refer to **Supplemental Table 2**). For all sequences, the transcripts per million (TPM) were used for absolute counts. Subsequent analysis used quantile normalization data that was log2 transformation and z-score normalization of expression in Python 3.11 and Pandas 1.5.3. Statistically significant differential expression using EBSeq-HMM was included in the RSEM distribution Leng2015-yz. EBSeq-HMM normalized differential expression was performed by quantile normalization. The significance for cutoff for differential expression was a False Discovery Rate (FDR) of <0.05. A CDS sequence for eGFP (Genbank: AAB02572.1) was included during transcript mapping to ensure that the RNA from sorted cells were indeed GFP-positive (**Supplemental Table 1**). For downstream analyses, alternate transcripts in the *D. melanogaster* genome were aggregated prior to gene level analysis using a custom script using Pandas 1.5.3 to aggregate data.

### Gene expression analyses

#### Co-expression clustering

Genome-wide co-expression clustering was performed using quantile normalized TPM counts per transcript, followed by log2 transformation and z-score transformation. Heatmaps were prepared in Python 3.11, Pandas 1.5.3, and Seaborn 0.12.2 clustermap function that implements the SciPy 1.10.0. linkage algorithm using Euclidean distance. Co-expression clustering was performed using the *clust* version 1.12.0 [49] on TPM counts for each transcript corresponding to genes identified as having statistically significant differential expression during the stages of border cell migration. *Clust* automatically performs quality control and normalization of TPM counts per gene and clusters expression patterns represent biological expectations [49].

#### Gene annotation enrichment and protein network analysis

Metascape was used for annotation enrichment analysis [50; 100] of genes identified in each of the *clust* co-expression clusters. Default analysis parameters were used and functional annotation results were further analyzed with customized visualizations in Python. Protein interaction networks derived from Metascape analysis of *clust* co-expression clusters were augmented with additional protein interactions and annotations from STRING [52], with a confidence cutoff score of 0.4 implemented as a plug-in in CytoScape version 3.9.1 [51]. Protein/gene nodes in networks without interactions, or those with a single interaction, were removed from further analysis. Metascape functional annotations augmented with STRING annotations were manually curated into a consistent set of functional terms reflective of the gene function based on gene summary, gene group and protein family, pathway, additional gene ontology (GO), and phenotype data from Flybase 2023_01 and Dmel Release 6.50 (“Annotations” column, **Supplemental Table 7**) [41]. For clarity, one annotation “keyword” was chosen for each protein in the interaction network (“Annotation Keyword” column, **Supplemental Table 7**). Protein/gene nodes were then colored based on annotation keyword to highlight regions of the network with similar functions as indicated. Additional manual curation was performed on immune and ribosome biogenesis related genes.

#### Immune and ribosome biogenesis pathway analysis

Differentially expressed genes were chosen based on Metascape enrichment for immune and ribosome related GO terms (including “innate immune response,” “immune effector process,” “ribonucleoprotein complex biogenesis,” “ribosomal small/large subunit complex biogenesis,” and “rRNA processing” (**Supplemental Tables 8-9**) [100]. To confirm biological functions for these chosen genes, we used FlyBase data [41] and primary literature to annotate each gene (“Annotation Notes” and “References” columns (**Supplemental Tables 8-9**). Based on this annotation, each gene was then either assigned a category or excluded from analysis (“Category” column; **Supplemental Tables 8-9**). Differentially expressed genes enriched for immune GO terms were then placed into either the Toll, Imd, JNK, or other signalings categories, or into a more general “immune response” category (**Figure 5**). For a more detailed view of each pathway and differential expression patterns for each gene, we combined expression and pathway information in **Supplemental Figure 2**.

#### Heatmaps of gene groups of interest

FlyBase gene numbers (FBgns) associated with GO IDs for oogenesis (GO:0048477), actin cytoskeleton (GO:0015629), and adhesion (GO:0007155, GO:0022610) were downloaded from FlyBase [41]. FBgn numbers for transcription factors were taken from the “PL FlyTF_trusted_TFs” list on FlyMine ([101; 102], https://www.flymine.org/flymine). Genes for epithelial-to-mesenchymal transition (EMT; GO:0001837) were obtained using the high rank orthologs from mouse EMT genes as determined by the DRSC Integrative Ortholog Prediction Tool (DIOPT) [103; 104] in addition to FBgns associated with the indicated GO IDs downloaded from FlyBase. “Border cell migration genes” were manually curated using primary literature to include genes likely to function in border cell migration (genes and references available in **Supplemental Table 4).** FBgns from Wang et al. [18] were taken from their Supplemental Table 1 (“Genes Enriched in Migratory Cells”). FBgns from Borghese et al. [17] were taken from their supplemental tables S1 and S4 (S1: “Genes significantly up-regulated; P<0.05, in wild type border cells [BCs] compared to follicle cells [WT BCs > FCs]”, S4: “Genes significantly [P<0.05] down-regulated in wild type border cells compared to follicle cells [WT BCs < FCs]”). FBgns were translated to FlyBase transcript numbers (FBtr) using FlyBase and expression of all significantly differentially expressed transcripts (EBseq-HMM FDR<0.05) was assessed.

#### Dresden Ovary Table RNA situ hybridization database analyses

Fluorescence *in situ* hybridization images for genes present in the dataset were accessed from the Dresden Ovary Table (DOT) website [60; 61]. The “gene search” and “table” functions were used individually to search for expression data for the 1,262 significantly differentially expressed genes identified in our analyses. Although the DOT provides developmentally staged images with annotated cell types, it was necessary to empirically validate these results. Therefore, images were manually assessed for each gene that had data for oogenesis stages 8-10 and the relevant cell types visible, including border cells, polar cells, centripetal cells, and posterior terminal cells (**Supplemental Table 10**). Expression data was manually curated for significantly differentially expressed genes except where images were not available for the given gene. In this case, the DOT annotation was used, which is denoted by use of italicized text in **Supplemental Table 10**. A homogenous RNA signal throughout the egg chamber was denoted as “ubiquitous signal.” Representative images were chosen for a subset of genes with strong RNA signal in border cells at stages 9 and/or 10 (**Figure 6**).

### Immunostaining and Microscopy

To analyze border cell migration, whole ovaries were dissected in Schneider’s *Drosophila* medium (Thermo Fisher Scientific) and fixed in 4% methanol-free formaldehyde (Polysciences) and 0.1 M potassium phosphate buffer (pH 7.4) for 10 minutes. To analyze migration, egg chambers were stained for E-cadherin (E-cad; 1:10 dilution, rat monoclonal DCAD2; Developmental Studies Hybridoma Bank [DSHB]), Singed (Sn; 1:25 dilution, mouse monoclonal Sn7C; DSHB), and DAPI to label nuclei (2.5 μg/mL; Millipore Sigma, cat. no. D954). Isotype-specific anti-mouse or anti-rat secondary antibodies conjugated to AlexaFluor–488 or –568 (Thermo Fisher Scientific) were used at a concentration of 1:400. All slides were mounted using Fluorsave Reagent mounting media (Millipore Sigma, cat. no. 345789). Migration defects were analyzed on an upright Zeiss AxioImager Z1. A primary antibody against GFP (1:200 dilution, mouse monoclonal 12E6; DSHB) was used to improve visualization of the *slbo*-mCD8:GFP pattern (**Figure 1A-C**). To visualize the *c306*-Gal4 expression pattern (**Figure 7A-C**), *c306*-Gal4; tsGAL80>UAS-LacZ egg chambers were stained with a primary antibody against beta-galactosidase (1:10 dilution, mouse monoclonal 40-1a; DSHB). All images were acquired on a Zeiss LSM 880 confocal microscope at the Kansas State University (KSU) College of Veterinary Medicine (CVM) Confocal Core using a 20x numerical aperture (NA) objective. All images were processed in ImageJ (FIJI).

### Statistical Methods, Graphs, and Figure Assembly

Bioinformatic analyses were performed in Python version 3.11, Pandas version 1.5.2, Numpy version 1.24.1, SciPy version 1.10.0, and Seaborn version 0.12.2 overlaid on Matplotlib version 3.6.3. Data tables were analyzed in Python and Pandas. Tables were exported to Microsoft Excel format and functions using OpenPyxl version 3.0.10. Additional graphs were assembled and statistical analyses were performed in GraphPad (version 7.04). Figures were assembled using Affinity Photo, Affinity Designer, and Adobe Illustrator. Three trials per RNAi line were used for the RNAi screen. Cutoff value for migration defect was determined by the background mean migration defect, calculated using average migration defect for the negative control (*c306*-Gal4; tsgal80/+; UAS-mCherry RNAi/+ [BDSC #35785]). To assess significant migration defects, the empirical rule for a normal distribution was used, where the migration defect cutoff is equal to the mean plus three times the standard deviation.

## Abbreviations

DOT: Dresden Ovary Table
EMT: epithelial-mesenchymal transition
FBgn: Flybase gene number
GO: gene ontology
PPI: protein-protein interactions
TPM: transcripts per million

## Declarations

### Ethics approval and consent to participate

Not applicable.

### Consent for publication

Not applicable.

### Availability of Data and Materials

Data and code used to generate figures will be available upon publication. RNA-sequencing data is available at NCBI Sequence Read Archive (SRA) BioProject ID (XXX, will be provided soon). Major results are available in the supplemental tables. All other data and materials will be made available on request.

### Author Contributions

Conceptualization – E.B., B.J.S.C.O., and J.A.M.; Data curation – J.R., A.T., and B.J.S.C.O.; Formal analysis – E.B., J.R., and B.J.S.C.O.; Funding acquisition – E.B., B.J.S.C.O. and J.A.M.; Investigation – E.B., J.R., A.T., P.M., and B.J.S.C.O.; Validation – E.B., J.R., and B.J.S.C.O.; Writing original draft – E.B., J.R., B.J.S.C.O., and J.A.M.; Writing – review & editing – E.B., J.R., A.T., P.M., B.J.S.C.O., and J.A.M.

## Acknowledgments

We would like to thank the Bloomington Drosophila Stock Center, the Harvard Transgenic RNAi Project, the Vienna Drosophila Resource Center, and Xiaobo Wang for providing flies, and the Developmental Studies Hybridoma Bank at the University of Iowa for providing antibodies used in this study. Sequencing was done by the University of Kansas (KU) Genome Sequencing Core, which is supported by the National Institute of General Medical Sciences (NIGMS) of the National Institutes of Health under award number P30GM145499. We also thank the Kansas State University College of Veterinary Medicine Confocal Core for use of the Zeiss LSM880 confocal. Thank you to Juliet Her for help with the Dresden Ovary Table RNA *in situ* hybridization image analysis, and to Yujun Chen and Rehan Khan for helpful comments on the manuscript.

## Supplementary Information

**Supplemental Figure 1.**
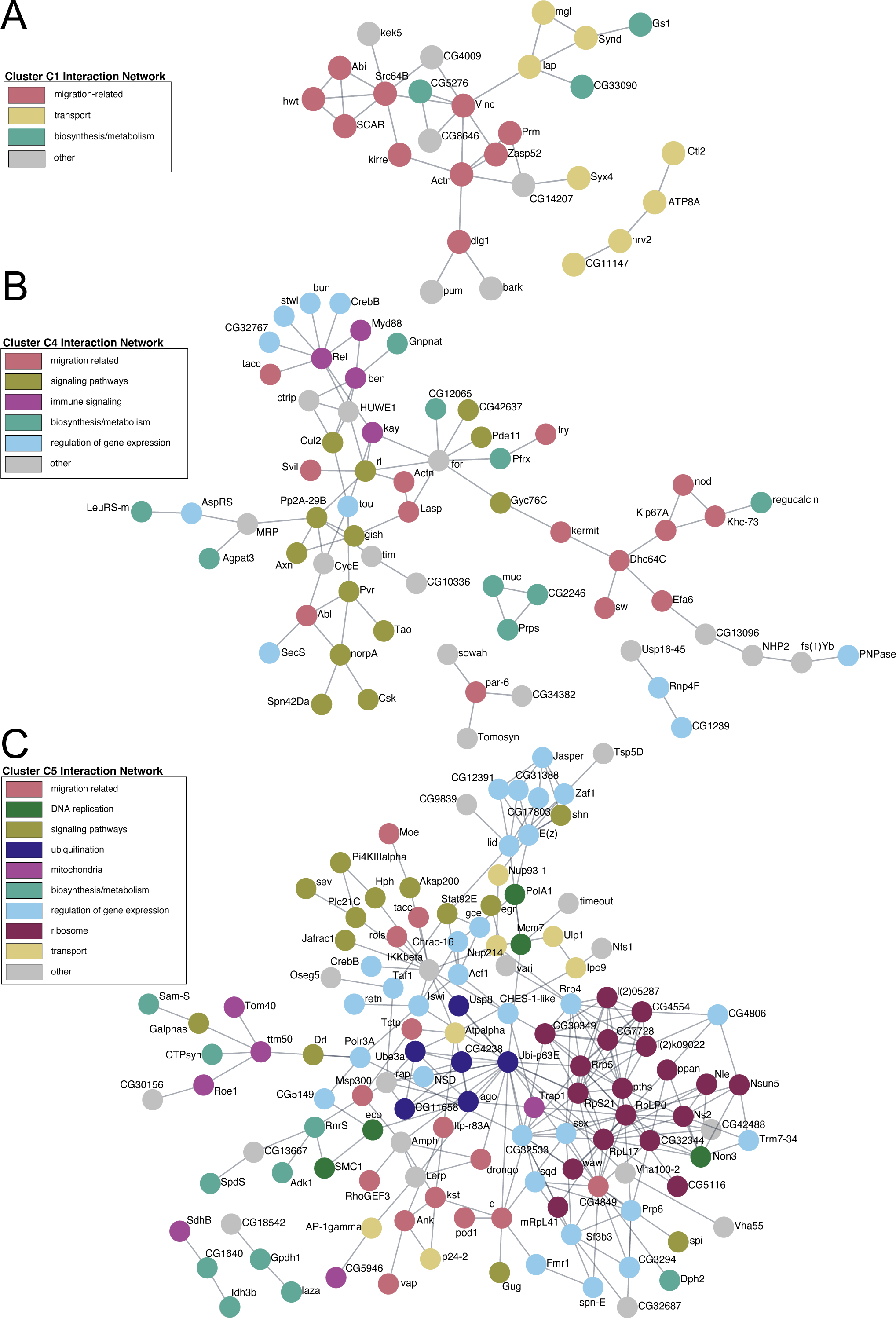
Protein-protein interaction networks for co-expression clusters C1, C4 and C5. (A-C) Physical PPI network analysis of genes in selected co-expression clusters C1 (A), C4 (B) and C5 (C). Individual small networks are found in each cluster in addition to the larger networks. (A) Network of C1 co-expressed genes with expression that increased during border cell migration. (B, C) Networks of C4 (B) and C5 (C) co-expressed genes with expression patterns that decrease during border cell migration. Functional annotation keywords (**Supplemental Table 7**) were used to assign color to proteins in the networks. (A) Co-expression cluster C1 forms a network with migration-related functions (pink). An individual node (top right) and network (lower right) have transport-related functions (mustard). (B) The co-expression cluster C4 network forms nodes with immune signaling (purple) and regulation of gene expression (light blue, upper left), biosynthesis/metabolism (cyan, center), and signaling pathways (green, center). (C) Co-expression cluster C5 forms a large node for ribosome function (magenta, right), nodes for regulation of gene expression (light blue, top, center, and lower right), and a node for signaling pathways (green, upper left).

**Supplemental Figure 2.**
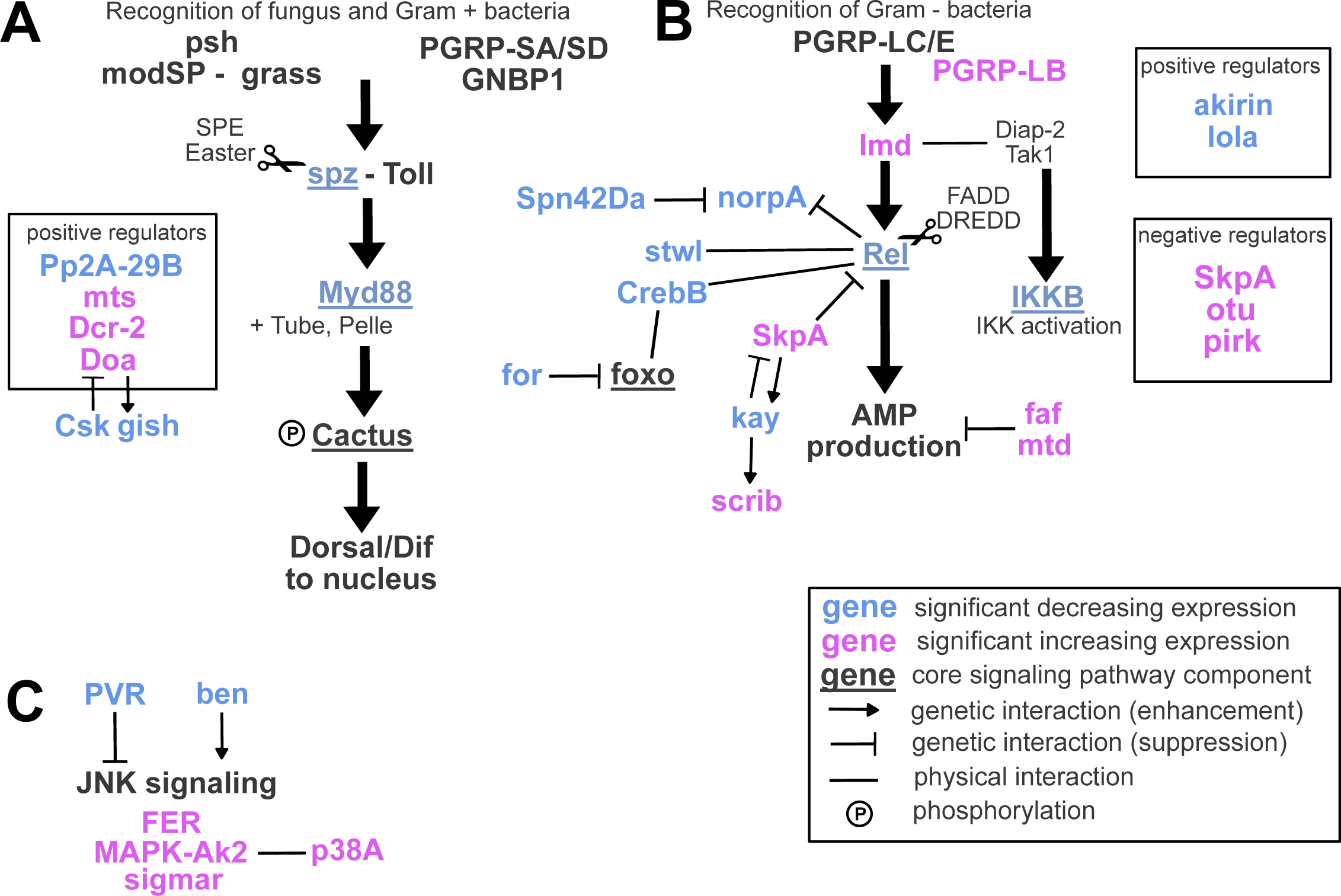
Toll, Imd, and JNK signaling pathway genes differentially expressed during border cell migration. Diagrams of key components and regulators of the Toll (A), Imd (B), and JNK (C) signaling pathways, some of which are differentially expressed during border cell migration. Genes that are differentially expressed in border cells are in bold text, while those that increase in expression are shown in magenta and those that decrease in expression are shown in blue. Genes denoted as core signaling pathway components on Flybase are underlined.

